# AraC binds the p75^NTR^ transmembrane domain to induce neurodegeneration in mature neurons

**DOI:** 10.1101/2021.12.01.470721

**Authors:** Vanessa Lopes-Rodrigues, Pia Boxy, Eunice Sim, Dong Ik Park, Josep Carbonell, Annika Andersson, Diana Fernández-Suárez, Anders Nykjær, Lilian Kisiswa

## Abstract

**Background:** Cytosine arabinoside (AraC) is one of the main therapeutic treatments for several types of cancer including acute myeloid leukaemia. However, after high dose AraC chemotherapy regime, patients develop severe neurotoxicity and neurodegeneration in the central nervous system leading to cerebellar ataxia, dysarthria, nystagmus, somnolence and drowsiness. AraC induces apoptosis in dividing cells, however, the mechanism by which it leads to neurite degeneration and cell death in mature neurons remains unclear. We hypothesized that the upregulation of the death receptor p75^NTR^ is responsible for AraC-mediated neurodegeneration and cell death in leukemia patients undergoing AraC treatment.

**Methods:** To determine the role of AraC-p75^NTR^ signalling in degeneration of mature cerebellar granule neurons, we used primary cultures from p75^NTR^ knockout and *p75^NTRCys259^* mice. Evaluation of neurodegeneration, cell death and p75^NTR^ signalling was done by immunohistochemistry and immunoblotting. To assess the direct interaction between AraC and p75^NTR^, we performed isothermal dose response-cellular thermal shift and AraTM assays as well as Homo-FRET anisotropy imaging.

**Results:** We show that AraC induces neurite degeneration and programmed cell death of mature cerebellar granule neurons in a p75^NTR^-dependent manner. Mechanistically, AraC binds to Proline 252 and Cysteine 256 of the p75^NTR^ transmembrane domain and selectively uncouples p75^NTR^ from the NF*κ*B survival pathway. This in turn, exacerbates the activation of the cell death/JNK pathway by recruitment of TRAF6 to p75^NTR^.

**Conclusion:** Our findings identify p75^NTR^ as a novel molecular target to develop treatments to counteract AraC-mediated neurodegeneration.

## Background

AraC (1-β-arabinofuranosylcytosine or Cytosine arabinoside) is the most effective chemotherapy agent used to treat patients with acute myeloid leukemia (AML), as well as other types of haematological cancers^1–5^. In order to achieve therapeutic efficacy of AraC, patients are subjected to a High Dose AraC (HIDAC) chemotherapy regime^1^. Although, HIDAC is efficient in treating AML, it leads to severe cerebellar neurotoxicity^6, 7^. It has been suggested that AraC elicits neurotoxicity by inducing programmed cell death in cerebellar neurons, in particular cerebellar granule neurons (CGNs)^8, 9^. However, this has only been shown for immature CGNs during development^8, 9^. Interestingly, AraC mediated neurotoxicity is age-dependent, with patients over 50 years old being more susceptible^3, 10, 11^. Generally, neuronal proliferation ceases after the developmental period, with exceptions of a few brain regions such as hippocampus and olfactory bulb^12, 13^. AraC has been shown to induce apoptosis in proliferating cells by inhibiting DNA synthesis^14, 15^ and DNA repair^16^. Therefore, it is unlikely that the mechanism by which AraC induces cerebellar neurodegeneration in adult patients is by targeting proliferating neurons. This warrants the elucidation of the mechanisms by which AraC induces degeneration of mature neurons.

The death receptor p75^NTR^ is highly expressed in the nervous system during development, but it is widely down-regulated in the adult brain^17^, with the exception of the cholinergic neurons in the basal forebrain^18^. However, low levels of expression of p75^NTR^ persist into adulthood in some areas of the central nervous system (CNS) such as cerebellum, septum, medulla and pons^19^. Moreover, a growing body of evidence have demonstrated an upregulation of p75^NTR^ under pathological conditions including cancer, brain injury and neurodegenerative diseases^20–, 23^. Interestingly, an increase in p75^NTR^ expression in the serum and peripheral blood of leukemia patients has been reported^24, 25^. Therefore, we asked whether p75^NTR^ could be responsible for mediating AraC-induced neurite degeneration and cell death in mature cerebellar neurons. Here, we show that AraC induces neurite degeneration and apoptosis in mature CGNs by binding to the transmembrane domain (TMD) of p75^NTR^, an interaction that is dependent on the Proline (Pro) 253 and Cysteine (Cys) 256 residues. Functionally, we show that AraC binding to the p75^NTR^ TMD uncouples p75^NTR^ from the NFkB survival pathway, resulting in the exacerbated activation of the cell death/JNK pathway in CGNs.

## Materials and methods

### Animals

Mice were housed in a 12-hour light/dark cycle and fed a standard chow diet. The transgenic mouse lines used were p75^NTR^ knockout (*p75^NTR-/-^* )^26^ and *p75^NTRCys259^* (knock-in mice carrying a substitution at amino acid position 259) ^27^ mice. Both transgenic mouse lines were maintained in a C57BL/6J background. Mice of both sexes were used for the experiments. All animal experiments were conducted in accordance with the National University of Singapore Institutional Animal Care, Use Committee and the Stockholm North Ethical Committee for Animal Research regulations and the Danish Animal Experiment Inspectorate Under the Ministry of Justice.

### Neuronal cultures

P7 mouse cerebella were dissected with removal of the meninges in ice-cold phosphate saline buffer (PBS). Whole cerebella were then digested with TrypLE™ Express (Gibco, 12604021) for CGNs extraction. CGNs were plated at a density of 40,000 cells per coverslip coated with poly-D-lysine (Sigma, P1524) in a 24-well plate (Thermo Scientific, 142475) in neurobasal medium (Gibco, 21103049) supplemented with 25 mM KCl, 1 mM Glutamax (Gibco, 35050061), 1X Pen/Strep (Sigma, P4333), 10 μM AraC (Sigma, C-6645), 1X B27 supplement (Gibco, 17504044). AraC was purchased from Sigma (C-6645), diluted in PBS and used at concentration of 10μM for elimination of glia cells in the neuronal cultures. For experimental conditions AraC was used at concentration of 500μM and 1000μM

#### (i) Neurite degeneration

To assess neurite degeneration, 4 DIV wild type (WT) neurons were treated for 24 or 48 hours with 500μM AraC. After treatment, CGNs were fixed for 10 minutes with ice-cold methanol. Cells were then permeabilized and blocked in 5% normal donkey serum and 0.3% Triton X-100 in PBS. Cells were incubated overnight at 4 °C with mouse anti-β-III tubulin (R&D Systems, MAB1195; 1:2500) and counterstained the following day with donkey anti-mouse Alexa Fluor 488 (Abcam, ab150105; 1:2000) and Hoechst (Sigma, B2261; 1:2000). Images were taken using a Confocal Microscope LSM 780 Zeiss Axio Observer fluorescence microscope. Images were captured from regions with well-separated neurites. NIH ImageJ software was used to threshold and binarize the images and remove all cell bodies, after which the Analyse Particles algorithm was applied to identify the area of fragments based on size (20– 10,000 pixels) and circularity (0.2-1.0). The degeneration index (DI) was then calculated as the ratio of the total area of detected neurite fragments over the total neurite area. In agreement with previous studies^28^ (, a DI of 0.2 or higher indicated neurite degeneration.

#### (ii) Cell death

Apoptosis was assessed in WT, *p75^NTR-/-^* and *p75^NTRCys259^* CGNs treated for 24 hours with either 500μM or 1000μM AraC starting at 4 DIV, Apoptotic cells were labelled using Click-iT plus TUNEL assay for in situ apoptosis detection kit (Thermoscientific, Cat: C10617) according to manufacturer instructions. Neurons were also stained for cleaved caspase 3, *β*-III tubulin and DAPI following protocol for immunocytochemistry explained below. For each experiment and treatment, neurons were culture in duplicates and at least 15 images were taken per coverslip with a Zeiss Axioplan confocal microscope. Quantification of number of cells positive for cleaved caspase 3 and TUNEL was performed using NIH ImageJ software.

#### (iii) Protein collection and immunoblotting

To collect protein for immunoblotting, WT neurons were cultured at high density (∼200,000 neurons per well) in a 48-well plate. 4 days after plating, cells were stimulated with 500μM AraC for 15, 30 and 60 minutes.

Protein samples were prepared for SDS-PAGE in SDS sample buffer and boiled at 95 °C for 10 min before electrophoresis on 12% gels. Proteins were transferred to PVDF membranes (Amersham). Membranes were blocked with 5% non-fat milk and incubated with primary antibodies.

The following primary antibodies were used at the indicated dilutions: rabbit anti-phospho Y515 TrkB (Abcam, Cat: ab131483, 1:500), goat anti-TrkB (R&D systems, Cat: AF1494, 1:500), rabbit anti-I*κ*B*α* (Santa Cruz, Cat: 9165, 1:500), rabbit anti-phospho-c-Jun (Thr91, cell signal, 2303, 1:1000), rabbit anti-c-Jun (cell signal, 9165, 1:1000) and mouse anti-GAPDH (Sigma, Cat: G8795, 1:1000). Immunoreactivity was visualized using appropriate HRP-conjugated secondary antibodies. Immunoblots were developed using the ECL Advance Western blotting detection kit (ThermoFisher scientific, Cat; 34095) and imaged using chemiluminescent western blot imaging system, Azure c300 (Azure biosystems). Image analysis and quantification of band intensities were done using NIH ImageJ software.

#### (iv) RhoA Assay

Protein was extracted from WT CGNs that were treated at 4 DIV with 500μM AraC for 30 minutes. RhoA activity was evaluated in total CGNs extracts using the RhoA G-Lisa kit (Cytoskeleton, Cat; BK124) following the manufacturer’s instructions. Equal amount of protein was used from each sample as determined by BCA protein Assay (ThermoFisher scientific, Cat; 23235).

#### (v) Proximity Ligation Assay (PLA)

4 DIV WT CGNs were treated with 500μM AraC for 10 min. After treatment, CGNs were fixed for 15 min in 4% paraformaldehyde (PFA)/4% sucrose, permeabilized, and blocked in 10% normal donkey serum and 0.3% Triton X-100 in PBS. Neurons were then incubated overnight at 4° C with anti-p75^NTR^ (Promega; G323A; 1:500) and anti-TRAF6 (Santa Cruz; sc-8490; 1:100) antibodies in PBS supplemented with 3% BSA. The Duolink In Situ Proximity Ligation kit (Sigma) was used as per the manufacturer’s instructions. Cells were imaged with an LSM Imager Z2 confocal microscope (Zeiss) to detect PLA signals. PLA puncta were quantified using NIH ImageJ software with the plugin synaptic puncta analyzer^29^.

### Immunocytochemistry

For immunocytochemistry, the cultures were fixed in 4% paraformaldehyde and 4% sucrose for 15 min and washed with PBS before blocking nonspecific binding and permeabilizing with blocking solution (5% donkey serum and 0.3% Triton X-100 in PBS) for 1 h at room temperature. Neurons were incubated overnight with the primary antibodies in diluted blocking solution to 1% donkey serum at 4°C. After washing with PBS, the neurons were incubated with the appropriate secondary antibodies.

The primary antibodies used in this study were: polyclonal anti-cleaved caspase 3 (Cell signal, Cat: 9761, 1:400), monoclonal anti–β-III tubulin (R&D systems, Cat: MAB1195, 1:10000), polyclonal anti-p75^NTR^ (Neuromics, Cat: GT15057, 1:500), polyclonal anti-TrkB (RnD systems, Cat: AF1494, 1:250), polyclonal anti-MAP2 (Abcam, Cat: ab5392, 1:2000) and polyclonal anti-P65NFkB (Santa Cruz, Cat: sc-372, 1:250).

Secondary antibodies were Alexa Fluor–conjugated anti-immunoglobulin from Life Technologies, Invitrogen, used at 1:1500 (donkey anti-rabbit IgG Alexa Fluor 555, Cat: A31572), donkey anti-mouse IgG Alexa Fluor 488, Cat: A21202, donkey anti-mouse IgG Alexa Fluor 555, Cat: A31570, donkey anti-goat IgG Alexa Fluor 488, Cat numbers: A11055, donkey anti-rabbit IgG Alexa Fluor 488, Cat: A32790, donkey anti-chicken IgG Alexa Fluor 647, Jackson, Cat: 703-496-155). Images were obtained using a Zeiss Axioplan confocal microscope. Quantification of number of cells positive for cleaved caspase 3 and TUNEL was counted using NIH ImageJ software.

### Cellular thermal shift assay (CETSA)

Cellular thermal shift assay was performed as previously described^30^. 293T HEK cells constitutively expressing p75^NTR^ protein were homogenized in a buffer containing 100 mM HEPES, 1 mM DTT, 10 mM MgCl2, protease inhibitor cocktail tablets (Roche, Cat: 11836153001) and phosphatase inhibitors (Roche, Cat: 04906837001). After 3 freeze-thaw cycles using liquid nitrogen, the lysate was centrifuged (20,000 g, 4 °C, 20 min) to collect supernatant. Protein concentration was measured using BCA assay. The same amount of protein lysate was aliquoted into different PCR tubes and simultaneously subjected to 6 different temperatures (37, 41, 45, 49, 53, 57 °C) for 3 min in the Veriti Thermal Cycler (Applied Biosystems). After 3 min cooling on ice, heat-treated protein lysate was centrifuged (20,000 g, 4 °C, 20 min). Supernatant was collected into new tubes, and samples were subjected to immunoblotting. Primary antibodies used were anti-p75^NTR^ (Promega, Cat: G323A, 1:300) and anti-GAPDH (Sigma, Cat: G8795, 1:1000).

### Isothermal dose response-cellular thermal shift assay (ITDR-CETSA)

The protein lysate from 293T HEK cells constitutively expressing p75^NTR^ protein were homogenized in the same buffer as described above for CETSA and incubated with different concentrations of AraC (0, 0.1, 1, 30, 100, 300, 500 and 1000μM) for 3 min at RT. The protein extracts were heated at a constant temperature of 37 or 53 °C for 3 min. After subsequent cooling on ice for 3 min, heat-treated lysate was centrifuged (20,000 g, 4°C, 20 min), followed by supernatant collection. Samples were then subjected to immunoblotting and probed for anti-p75^NTR^ (Promega, Cat: G323A, 1:300) and anti-GAPDH (Sigma, Cat: G8795, 1:1000). Densitometry data acquired from 37 °C ITDR-CETSA were used as non-denaturing controls to normalize those from ITDR-CETSA conducted at 53 °C. The densitometry was done on the immunoblot bands and thermal shift curve fitting was performed using Boltzmann sigmoidal equation (GraphPad Software, Inc., La Jolla, CA, USA)

### Homo-FRET anisotropy imaging

COS-7 were cultured under standard conditions in DMEM supplemented with 10% fetal bovine serum, 100 units/ml penicillin, 100 mg/ml streptomycin and 2.5 mM glutamate. Cells were transiently transfected with a rat p75^NTR^-EGFP* fusion constructs^31^ using FuGENE transfection reagents (Fisher scientific, Cat: PRE2311). EGFP* consist of a monomeric A207K EGFP mutant. 24 hours later anisotropy imaging was done as previously described^27, 32, 33^. Changes in anisotropy were expressed as fold change at each time point in comparison to the mean of 6 time points obtained prior to addition of vehicle or 500μM AraC. Images were acquired using Nikon Ti-E based live cell epi-fluorescence microscope and MetaMorph software and analyzed using MatLab from Mathworks.

### AraTM assay

AraTM assay was used to asses conformation changes and binding strength in a pair interaction of TMDs^34, 35^, as previously described^33^. Briefly, TMD cDNAs of the human p75^NTR^ (NLIPVYCSILAAVVVGLVAYIAFKRW) and TrkB (SVYAVVVIASVVGFCLLVMLFLL) were subcloned into AraTM chimera plasmid in between the KpnI and SacI restriction sites. AS19 LPS-negative *E.Coli* cells were transformed with each of the above mentioned plasmids together with a GFP reporter plasmid. The selected colonies were grown overnight for 18 hours in the shaker at 37°C in Lysogeny Broth (LB) containing 50 μg/ml, Spectinomycin and 100 μg/ml. Then the culture was then diluted 1:100 in fresh LB medium and allowed to grow till optical density (OD) 630 reached between 0.2 and 0.5 after which 1mM IPTG was added to induce the expression of the p75^NTR^ TMD–AraTM chimera or TrkB TMD-AraTM chimera. 100 μl of the culture per well were dispensed in black-rim clear bottom 96-well plates (Corning, cat: 3631) previously plated with serial concentration (30, 100, 300, 500 and 100μM) of AraC. The plates were then incubated at 38^0^C on shaking for 4 hours to allow IPTG-induced expression of the TM-AraTM chimera. The plates were then centrifuged at 4000 rpm for 10 min at room temperature to pellet bacteria. LB media was aspirated and replaced with 100 μl of Phosphate Buffered Saline (PBS), and bacteria cells were resuspended by vigorous shaking for 10 min. GFP signal was measured in each well (excitation 475nm, emission 509 nm) and bacterial density was determined by measurement of OD630 in a microplate plate reader (BioTek).

### Statistical analysis

Data are expressed as mean and standard errors of the mean. No statistical methods were used to predetermine sample sizes but our sample sizes are similar to those generally used in the field. Following normality test and homogeneity variance (*F*-test or Kolmogorov-Smirnov test with Dallal-Wilkinson-Lilliefor P value), group comparison was made using an unpaired student *t*-test, one-way or two-way ANOVA as appropriate followed by Bonferroni post-hoc test for normally distributed data. Differences were considered significant for *P* < 0.05. The experiments were not randomized.

## Results

### AraC induces neurite degeneration and apoptosis in mature CGNs

Apoptosis of immature CGNs associated with AraC is well-documented ^8, 9, 36–38^. However, it is known that adult cancer patients under AraC medication also develop cerebellar neurotoxicity resulting in neurodegeneration^6, 7^ despite lack of neurogenesis and proliferation in adult cerebellum. First, we aimed to confirm that AraC induces degeneration and apoptosis in mature CGNs. To obtain mature neurons for the experiments, we performed P7 cerebellar cultures, initially containing a mixture of glial cells and immature CGNs. To eliminate the proliferating glial cells from the culture, we cultured the cells for 4 days in media containing a low concentration of AraC (10μM: Fig 1a), obtaining an enriched CGN culture. The enriched CGNs cultures after 4 DIV (days *in vitro*) contain mature-like neurons that present high expression of mature markers such as MEF2 and Zic2 and low levels of immature markers such as Math1 or TAG^39^. On the 4 DIV, neurons were left untreated (control) or treated with either 500μM AraC for 24 or 48 hours (Fig 1a). Quantification of the neurite degeneration index showed an increase in neurite degeneration in neurons treated with 500μM AraC for 24 hours compared to untreated neurons (Fig 1b-c). Longer (48 hours) exposure of neurons to 500μM AraC worsened the neurite degeneration (Fig 1b&d). Moreover, cells treated with either 500μM or 1000μM AraC for 24 hours showed an increase in the number of cleaved caspase 3 positive neurons (Fig 1e-f). In agreement with this, analysis of the number of TUNEL analysis showed approximately 2-fold increase in apoptotic activity in CGNs after either 500μM or 1000μM AraC-treatment (Fig1g-h), suggesting that at the lower concentration the plateau on the apoptotic effect of the drug has been reached already. Therefore, these results are in agreement with previous studies^8, 9, 37^, indicating that AraC induce neurite degeneration and apoptosis in CGNs

**Figure 1:**
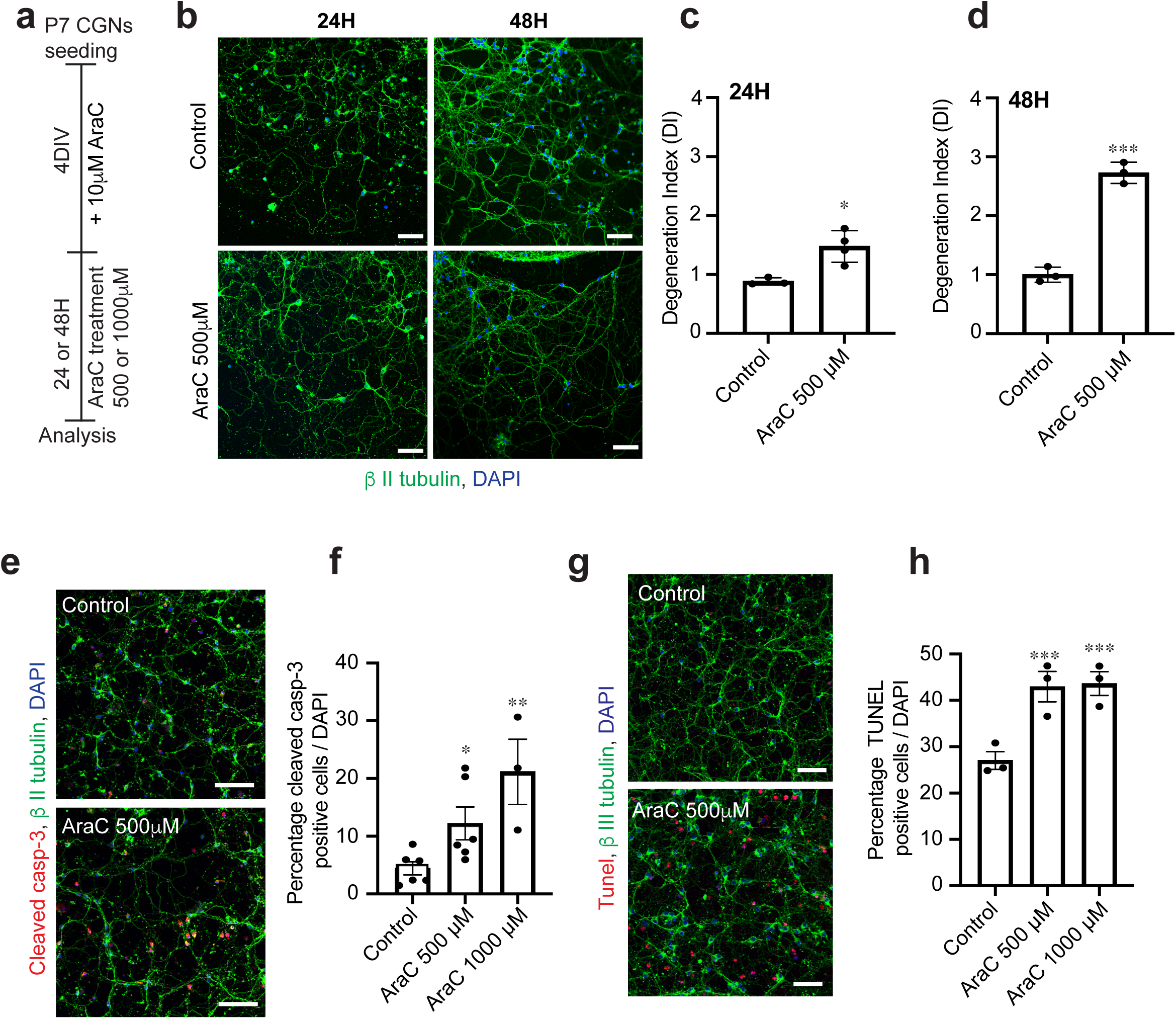
AraC induces neurite degeneration and apoptosis in mature CGNs. **(A)** A schematic drawing of the cell culture procedure. (**B**) Image of representative wild type P7 CGNs cultured for 4DIV that were either untreated (control) or treated with 500μM AraC for 24 or 48 h and stained for β III tubulin (green) and counterstained with DAPI (blue). Scale bars, 50 μm. (**C-D**) Quantification of degeneration index in control or CGNs treated with 500μM AraC for 24 h (**C**) and 48 h (**D**) CGNs cultures (total of 25 images per condition were quantified). Mean ± s.e.m. of data from four separate cultures (\**P* < 0.05 and \*\*\**P* < 0.001 compared to control, Unpaired Student T-test) is shown. (**E**) Image of representative wild type P7 CGNs cultured for 4DIV that were either untreated (control) or treated with 500μM AraC for 24 h and stained for cleaved caspase 3 (red), anti-β III tubulin (green) and counterstained with DAPI (blue). Scale bars, 50 μm. (**F**) Quantification of percentage cleaved caspase 3 positive neurons in control or CGNs treated with AraC (500μM or 1000μM for 24 h (total of 80 images per condition were counted). Mean ± s.e.m. of data from four separate cultures (\**P* < 0.05 and \*\**P* < 0.01 compared to control, one-way ANOVA followed by Bonferroni post-hoc test) is shown. (**G**) Photomicrographs of representative wild type P7 CGNs cultured for 4DIV that were either untreated (control) or treated with 500μM AraC for 24 h and stained for TUNEL (green), with anti-β III tubulin (red) and counterstained with DAPI (blue). Scale bars, 50 μm. (**H**) Quantification of percentage TUNEL positive neurons in untreated (control) or CGNs treated with AraC (500μM or 1000μM for 24 h (total of 60 images per condition were counted). Mean ± s.e.m. of data from three separate cultures, \*\*\**P* < 0.001 compared to control, one-way ANOVA followed by Bonferroni post-hoc test) is shown. Abbreviations: AraC, 1-beta-D-arabinofuranosyl-cytosine cytarabine; CGNs, cerebellar granule neurons; DIV, days in vitro

### p75^NTR^ death receptor is required for AraC-induced neurodegeneration and apoptosis in mature CGNs

We and others have shown that p75^NTR^ death receptor plays an important role in CGNs apoptosis during development^40–44^. Although p75^NTR^ is abundantly expressed in several types of developing neurons, its expression is negligible in the majority of mature neurons ^19, 45^. However, some areas of the adult brain, such as the cerebellum, contain low levels of p75^NTR^, and interestingly, it has also been observed that upon injury and neurodegeneration, p75^NTR^ expression is upregulated in the CNS^20, 22, 46^ . We therefore asked whether the cerebellum is especially vulnerable to AraC-induced neurodegeneration due to the presence of p75^NTR^ in mature cerebellar neurons. First, we confirmed the expression of p75^NTR^ in mature CGNs, as well as the presence of TrkB (Fig 2a), whose activity can be modulated by p75^NTR 47^. Next, we tested whether AraC-mediated neuronal death requires p75^NTR^. Wild type (WT) and p75^NTR^ knockout (*p75^NTR-/-^*) CGNs were treated at 4 DIV with AraC for 24 hours and then apoptotic activity was evaluated using cleaved caspase 3 and TUNEL assays. As expected, the number of WT CGNs positive for cleaved caspase 3 (Fig 2 b-c) and TUNEL (Fig 2 b&d) increased 2-fold after AraC treatment. Surprisingly, *p75^NTR-/-^* neurons treated with AraC did not show an increase in apoptotic activity (Fig 2 b-d). Altogether, these data suggest that AraC-mediated neuronal death requires p75^NTR^ death receptor.

**Figure 2:**
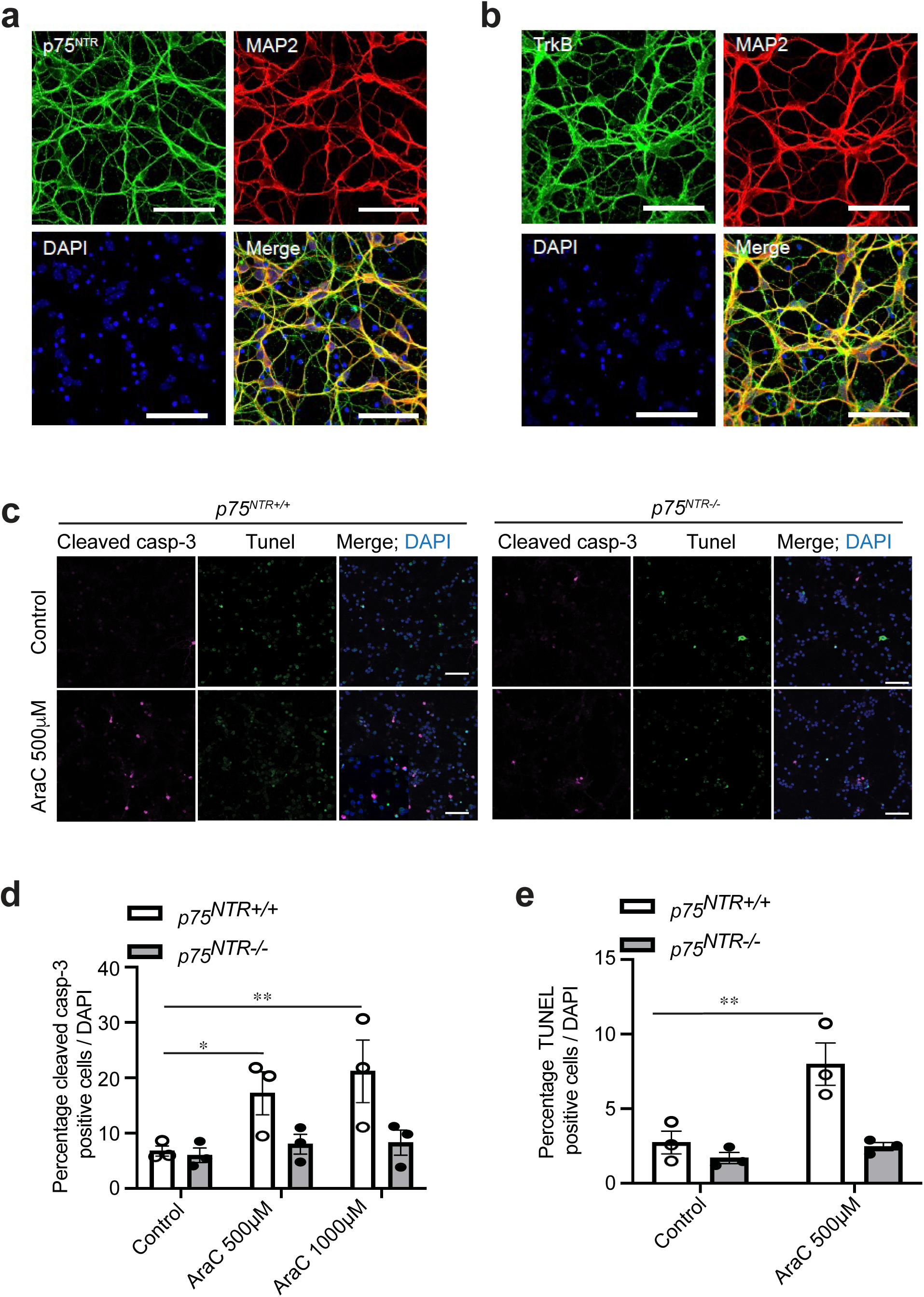
AraC requires p75^NTR^ to induce apoptosis in mature CGNs. (**A**) Representative micrographs of wild type P7 CGNs cultured for 4DIV and double stained with anti-p75^NTR^ together with anti-*β* III tubulin and counterstained with DAPI. Scale bars, 50 μm. (**B**) Representative micrographs of wild type P7 CGNs cultured for 4DIV and double stained with anti-TrkB (lower panel) together with anti-*β* III tubulin and counterstained with DAPI. Scale bars, 50 μm. **(C**) Image of representative P7 *p75^NTR+/+^ and p75^NTR-/-^* CGNs cultured for 4DIV, untreated (control) or treated with 500μM AraC for 24 h and stained for anti-cleaved caspase 3 (magenta), TUNEL (green) and counterstained with DAPI (blue). Scale bars, 50 μm. (**D-E**) Quantification of percentage cleaved caspase 3 positive (**D**) and TUNEL positive (**E**) *p75^NTR+/+^ and p75^NTR-/-^* neurons in untreated (control) or *p75^NTR+/+^ and p75^NTR-/-^* CGNs treated with AraC (500μM or 1000μM for 24 h (total of 80 images per condition were counted)). Mean ± s.e.m. of data from four separate cultures, \**P* < 0.05 and \*\**P* < 0.01 compared to control, one-way ANOVA followed by Bonferroni post-hoc test) is shown. Abbreviations: AraC, 1-beta-D-arabinofuranosyl-cytosine cytarabine; CGNs, cerebellar granule neurons; DIV, days in vitro

### AraC interacts directly with p75^NTR^

Although our data show that in the absences of p75^NTR^, AraC is unable to induce apoptosis in mature neurons, it was still unclear whether this effect was a result of a direct interaction between AraC and p75^NTR^ or an indirect effect. To assess this, we used the cellular thermal shift assay (CETSA). CETSA monitors protein-drugs interaction by assessing ligand-induced changes in the thermal stability of the protein of interest^48^. First, using protein lysate from 293T HEK cells that has constitutive expression of p75^NTR^, we generated the p75^NTR^ melting curve and found that p75^NTR^ starts to melt at 53°C (Data not shown). Then, using the lysate from p75^NTR^ expression HEK cells, we performed isothermal dose response (ITDR)-CETSA to assess the p75^NTR^ protein thermal stability at 37°C and 53°C after addition of different concentrations of AraC. While addition of AraC shifted thermal stability of p75^NTR^ in general, the higher concentrations (100-1000μM) of AraC led to a marked increase in p75^NTR^ destabilization at 53°C (Fig 3a-b) suggesting that AraC binds directly to p75^NTR^ resulting in thermal instability of the protein. Next, we sort to confirm that AraC could still bind to p75^NTR^ in intact cells using homo-FRET anisotropy assay. COS-7 cells were transfected with plasmid expressing a full-length rat p75^NTR^-EGFP* fusion^31^ and 24 hours later the changes in anisotropy levels after treatment with DMSO (control) or AraC were recorded over time. Treatment with AraC induced oscillation of p75^NTR^ anisotropy at the cells membrane (Fig 3c-e) resulting in a positive net change of the integrated peak area over 15 min treatment compared with vehicle (Fig 3d-e). Interestingly, the frequency of the anisotropy oscillation after AraC treatment was similar in the anisotropy changes above or under control basal line (DMSO average anisotropy values) (Fig 3f). Together, these data demonstrate that AraC binds directly to the p75^NTR^.

**Figure 3:**
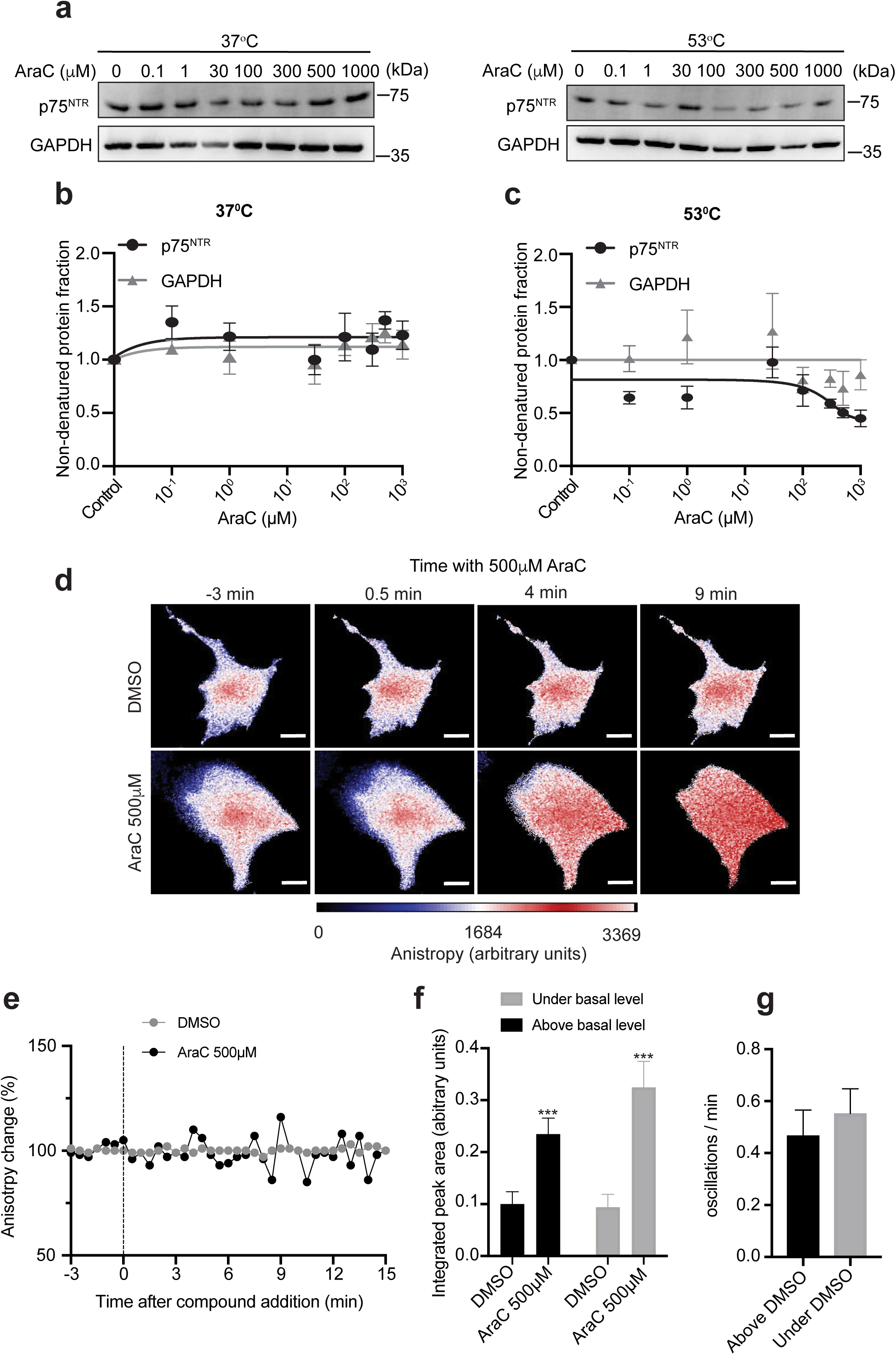
AraC binds to p75^NTR^. (**A**) Representative immunoblot of CETSA for AraC dose response on protein lysate extracted from HEK293 cells constitutively overexpressing p75^NTR^. The lysate was heated to 37°C and 53°C. (**B-C**) Quantification of p75^NTR^ and GAPDH protein levels at non-denaturing 37 °C upon binding AraC (**B**) and p75^NTR^ and GAPDH protein leves at 53 °C normalized to that of respective non-denaturing 37°C controls (**C**). GAPDH thermal stability was shown as a control for AraC specificity to p75^NTR^. Mean ± s.e.m. of data from five and three separate cultures is shown for p75^NTR^ and GAPDH, respectively. (**D**) Live cell homo-FRET anisotropy of p75^NTR^ in COS-7 cells in response to AraC. Shown are representative time lapse images before (-3 min) and after (0.5, 4 and 9 min) addition of 500 µM AraC or DMSO control. Scale bars, 5μm. (**E**) Live cell homo-FRET anisotropy of p75^NTR^ in COS-7 cells in response to AraC. Shown are representative traces of average anisotropy change after addition of AraC (500 µM at 0 min) or vehicle in cells expressing wild-type rat p75^NTR^. (**F**) Integrated peak area of live cell homo-FRET anisotropy of p75^NTR^ in COS-7 cells in response to AraC (considering area under and above y = 1). Results are plotted as means ± SD (N = 7). ***p < 0.001; Student T-test. (**G**) Oscillations/min of live cell homo-FRET anisotropy of p75^NTR^ in COS7 cells in response to AraC (considering the vehicle oscillations as threshold). Results are plotted as means ± SD (N = 7). Abbreviations: CETSA, the cellular thermal shift assay, AraTM, Ara transmembrane; FRET, Förster Resonance Energy Transfer; COS-7 - African green monkey kidney fibroblast-like cell line.

### AraC induces apoptosis in mature CGNs by binding to the Pro 253 and Cys 256 residues of the TMD of p75^NTR^

It was recently reported that a small molecule (NSC49652) binds to the p75^NTR^ TMD and induce cell death both in neurons and cancer cells^33^. We therefore, speculated that AraC might be binding to the transmembrane domain (TMD) of p75^NTR^. To this end, we evaluated whether AraC could bind to the TMD of p75^NTR^ using an AraTM assay where the bacteria expressed only the TMD of p75^NTR^. To assess the specificity of AraC binding to p75^NTR^ TMD, we used TMD from a different receptor, namely TrkB. We found that AraC preferentially binds to p75^NTR^ TMD (Fig 4a) although there was some minor binding to TrkB TMD (Fig 4a).

**Figure 4:**
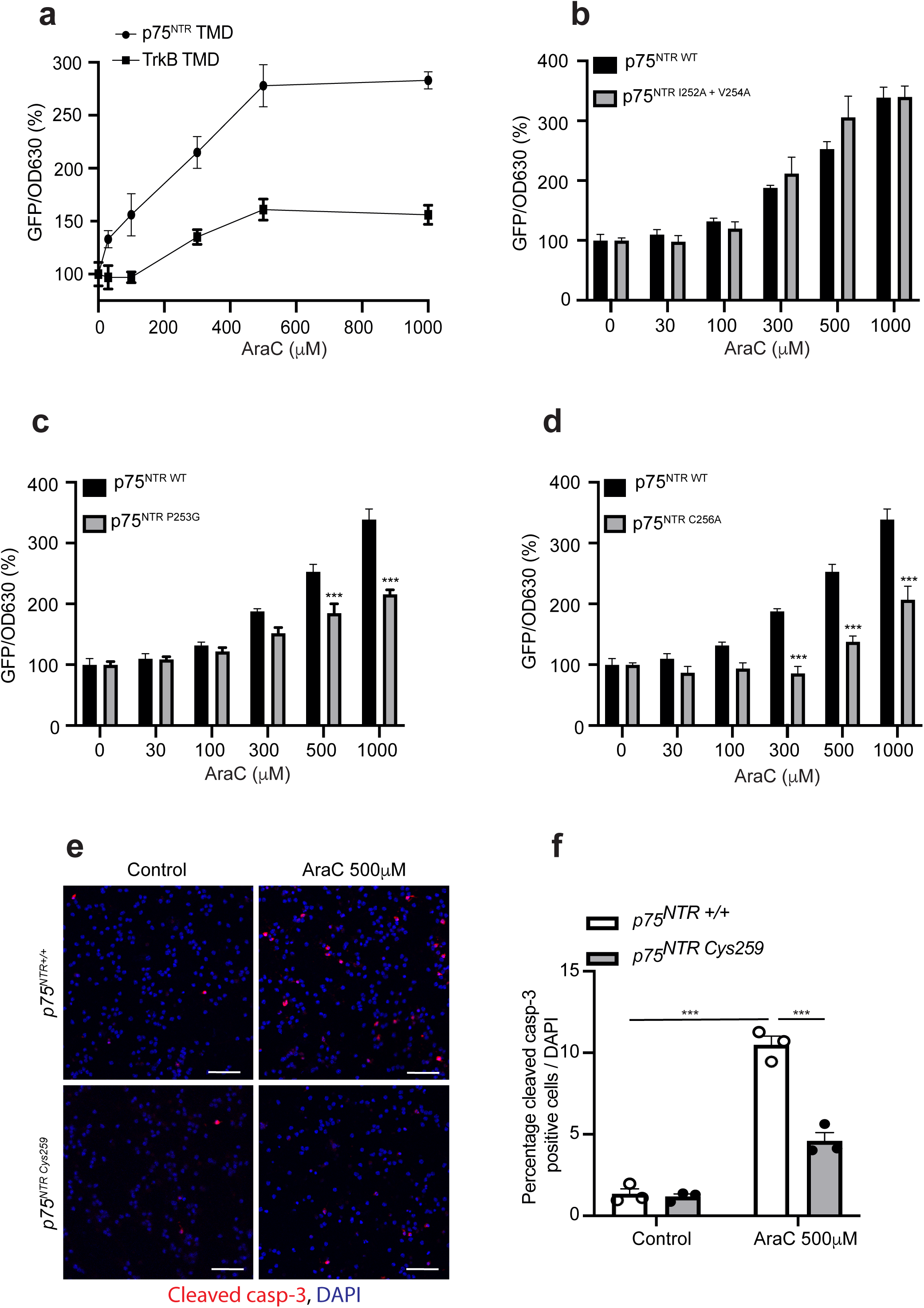
AraC induces programmed cell death by binding to Pro253 and Cys256 residues in transmembrane domain of p75^NTR^. (**A**) Dose response of AraC in the AraTM assay of p75^NTR^ and TrkB. Results are plotted as means ± SD (N = 3). **(B-D)** Comparison of human wild-type p75^NTR^ TMD with p75^NTR^ TMD with I252A plus V254A (**B**) or P253G (**C**) or C256A (**D**) mutants in the AraTM assay in response to increasing doses of AraC. The GFP over OD630 signal without any drug added was set at 100%. Results are plotted as means ± SD (N = 3). Significance was calculated using the 2-way ANOVA followed by Bonferroni post-hoc test where ** p< 0.01 and ***p < 0.001 (**E**) Representative images of wild type or C259A mutant P7 CGNs cultured for 4DIV that either were untreated (control) or treated with 500μM AraC for 24 h and stained for cleaved caspase 3 (red) and counterstained with DAPI (blue). Scale bars, 50 μm. (**F**) Quantification of percentage cleaved caspase 3 positive neurons in untreated (control) or wild type or C259A mutant CGNs treated with AraC (500μM) for 24 h (total of 60 images per condition were counted). Mean ± s.e.m. of data from four separate cultures, \*\*\**P* < 0.001 compared to control, one-way ANOVA followed by Bonferroni post-hoc test) is shown. Abbreviations: AraTM, Ara transmembrane; TMD, transmembrane domain.

After confirming the binding of AraC to the p75^NTR^ TMD, we sought to determine which residues facilitated this binding. The residues of human p75^NTR^ TMD are as follows NLIPVYCSILAAVVVGLVAYIAFKRW, starting at residue 250. Using a plasmid carrying human p75^NTR^ TMD, we started by mutating Isoleucine 252 and Valine 254 residues to Alanine (I252A and V254A, respectively), located at the beginning of p75^NTR^ TMD. Both bacteria transfected with human wild type p75^NTR^ TMD and p75^NTR^ TMD carrying the double mutants, I252A and V254A responded similar to AraC treatment (Fig 4b) suggesting that these two residues are not required for the binding of AraC to p75^NTR^ TMD. Next, we tested human p75^NTR^ TMD carrying mutation P253G, where Pro 253 was replaced with glycine. Upon AraC treatment, the bacteria carrying the P253G mutation had a lower percentage GFP/OD630 compared to bacteria carrying wild type p75^NTR^ TMD (Fig 4c). This result suggests that the Pro 253 is required for AraC binding to p75^NTR^ TMD. We then tested a point mutation, in which the Cys 256 was replaced with an alanine (C256A mutant). Similar to P253G, the C256A mutation had a lower percentage GFP/OD630 compared to wild type p75^NTR^ TMD upon AraC treatment (Fig 4d). Interestingly the response of the C256A mutant was already significantly diminished at lower concentration (300μM) of AraC, while the P253G mutant started to show alterations in the response at 500μM of AraC (Fig 4c-d), suggesting that the Cys 256 is crucial for AraC binding. Taken together, these data demonstrate that Pro 253 and Cys 256 are required for the binding of AraC to p75^NTR^ TMD.

Since the above-mentioned experiments were done in bacteria expressing only the TMD of p75^NTR^ and in transfected settings, we aimed to confirm the interaction of AraC with the TMD of the endogenous p75^NTR^ in mature neurons. To this end, we used CGNs from WT or C259A mice, treated the cells at 4 DIV with AraC and assessed for apoptosis 24 hours later. As expected from our previous results, treatment of wild type CGNs with AraC increased the percentage of cleaved caspase 3 positive cells in the cultures (Fig 4e-f). On the other hand, although C259A CGNs treated with AraC also showed an increase in the number of cleaved caspase 3 positive neurons, the number of apoptotic cells was 50% lower than in the WT neurons (Fig 4e-f). These results indicate that C259 is involved in the binding of AraC to the TMD of p75^NTR^, but other residues are necessary since that mutation does not completely block the apoptotic effect induced by the drug. All together, these data suggest that: (i) AraC binds to the TMD of p75^NTR^; (ii) the Pro253 and Cys256 (equivalent to mouse Cys259) residues are required for the binding of AraC to the TMD of p75^NTR^; (iii) AraC induces degeneration in CGNs by binding to the TMD of p75^NTR^.

### Binding of AraC to p75^NTR^ TMD does not affect p75^NTR^-dependent TrkB and RhoA signalling

Next, we dissected the signalling mechanism that AraC/p75^NTR^ employs to elicit cell death of mature CGNs. P75^NTR^ is known to engage several signalling pathways by interacting with other receptors and recruiting a number of adaptor proteins to its death domain, which lacks enzymatic activity^23, 42, 43, 49–53^. Moreover, p75^NTR^ has been suggested to increase the affinity of neurotrophins for Trk receptors^47, 54, 55^. To dissect the signalling mechanism that AraC/p75^NTR^ employs to elicit cell death of mature CGNs, we first evaluated whether the binding of AraC to p75^NTR^ TMD interferes with the activation of the TrkB signalling pathway. Upon activation by neurotrophins or pharmacological drugs, TrkB is phosphorylated on several residues including tyrosine 515 ^56, 57^. Phosphorylation of tyrosine 515 regulates protein kinase B (AKT) that plays an important role is cell survival^58^. We therefore evaluated the phosphorylation of TrkB, particularly on tyrosine 515 observing no changes upon AraC treatment (Fig 5a-c). This result suggests that the binding of AraC to p75^NTR^ TMD does not interfere with p75^NTR^-dependent TrkB activity. We then asked whether the AraC/p75^NTR^ interaction affects the signalling pathways (RhoA, JNK and NF*κ*B pathways) that are downstream to p75^NTR 23, 53^. We first assessed the AraC/p75^NTR^ interaction might modulate p75^NTR^-dependent RhoA activity in these neurons. However, CGNs treated at 4 DIV with AraC did not show any alteration in the levels of RhoA activity (Fig 5d).

**Figure 5.**
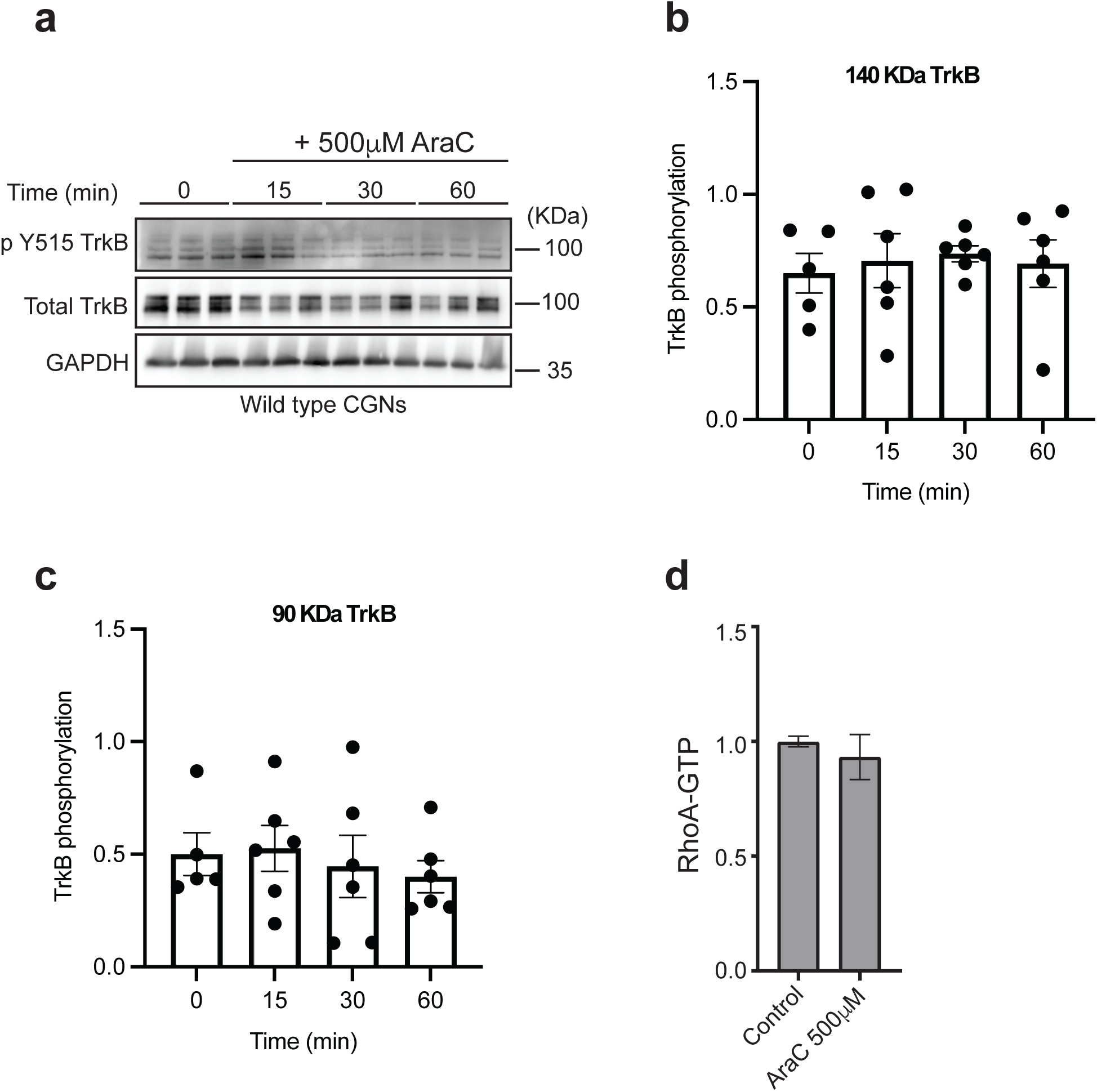
Binding of AraC to p75^NTR^ does not affect TrkB autophosphorylation or activation of RhoA pathway. (**A**) Representative western blots probed with phospho-TrkB (Y515), total TrkB and GAPDH of lysates of wild type P7 CGNs grown for 4 days prior to 15-, 30- or 60-minutes treatment with 500μM AraC. (**B-C**) Quantification of phosphorylation of the 140 KDa TrkB (**B**) and 90 KDa TrkB (**C**) in total lysate of untreated wildtype P7 CGNs or neurons treated with 500μM AraC for 15, 30 or 60 minutes. Mean ± sem of densitometry from 6 separate experiments is shown. (**D**) Analysis of RhoA-GTP levels in cerebellar extracts prepared from P7 wild type. Mean ± sem of data from 3 experiments is shown is shown.

### AraC selectively uncouples p75^NTR^ from NFkB signalling pathway

We previously showed that activation of p75^NTR^-dependent NFkB signalling pathway is crucial for survival of CGNs^43^. Therefore, we evaluated whether AraC treatment induces uncoupling of p75^NTR^ from the NFkB signalling pathway. p75-mediated activation of the NFkB pathway leads to the phosphorylation and subsequent degradation of IκBα (nuclear factor of kappa light polypeptide gene enhancer in B-cells inhibitor, alpha), which does not cover the nuclear localization signal of the cytosolic P65NF*k*B anymore leading to its translocation to the nucleus^43, 51^. Neurons treated with AraC for 30 minutes at 4 DIV showed lower P65NF*κ*B immunoreactivity in the nucleus compared to control (Fig 6a-b), suggesting that P65NF*κ*B remains bound to I*κ*B*a* in the cytosol. Therefore, we evaluated the degradation of I*κ*B*α* in untreated (naïve) cells and after treatment with 500μM AraC. We observed a decrease in I*k*B*a* degradation after 30 and 60 min of AraC treatment (Fig 6c-d).

**Figure 6.**
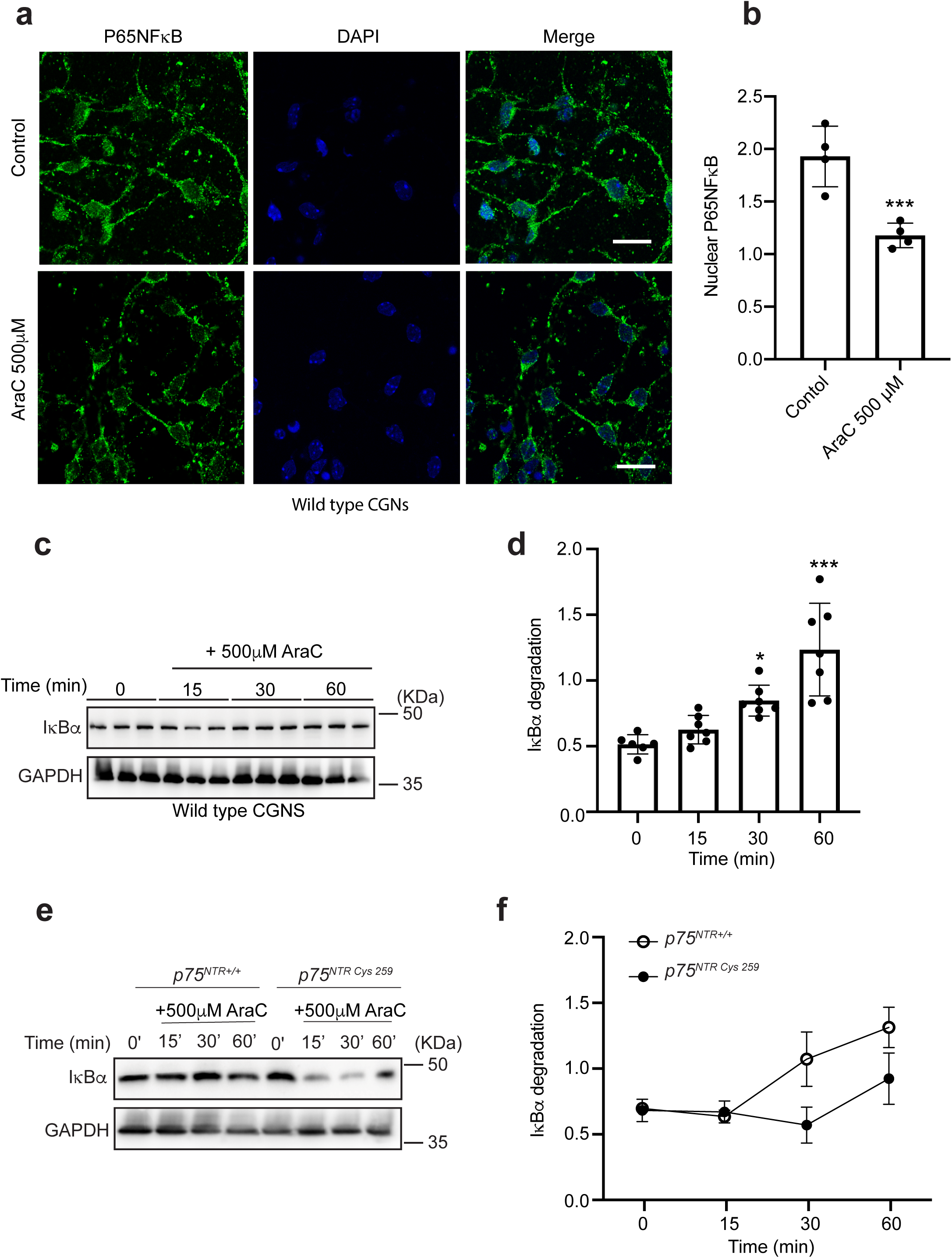
AraC selectively uncouples p75^NTR^ from NF*κ*B signalling pathway. (A) Representative images of wild type P7 CGNs cultured for 4 days, untreated (control) or treated with 500μM AraC for 60 minutes, fixed, stained for P65NF*κ*B (green) and counterstained with DAPI (blue). Scale bar, 50μM. (**B**) Quantification of the P65NF*κ*B nuclear translocation in untreated (control) or neurons treated with 500μM AraC for 60 minutes. Mean ± s.e.m. of data from four separate cultures, \*\*\**P* < 0.001 compared to control, Unpaired Student T-test) is shown. (**C**) Representative western blots probed with I*κ*B*α* and GAPDH of lysate of wild type P7 CGNs grown for 4 days prior to stimulation with 500μM AraC for 15, 30 or 60 minutes. (**D**) Quantification of I*κ*B*α* degradation in total lysate of untreated wildtype P7 CGNs or neurons treated with 500μM AraC for 15, 30 or 60 minutes. Mean ± sem of densitometry from 6 experiments (***P < 0.001; one-way ANOVA followed by Bonferroni test) is shown. (**E**) Representative western blots probed with I*κ*B*α* and GAPDH of lysate of wild type and C259A mutant P7 CGNs grown for 4 days prior to stimulation with 500μM AraC for 15, 30 or 60 minutes. (**F**) Quantification of I*κ*B*α* degradation in total lysate of untreated wildtype and C259A mutant P7 CGNs or neurons treated with 500μM AraC for 15, 30 or 60 minutes. Mean ± sem of densitometry from 3 experiments is shown.

Finally, we evaluated I*κ*B*α* degradation in WT and C259A neurons. As expected, WT neurons had an accumulation of I*κ*B*α* in the cytosol, while C259A neurons had a less pronounced I*κ*B*α* accumulation after AraC treatment (Fig 6e-f).

Taken together, these data indicate that the binding of AraC to the p75^NTR^ TMD leads to uncoupling of p75^NTR^ from the NF*κ*B signalling pathway resulting in the accumulation of IkB*α* in the cytosol and less translocation of P65NF*κ*B to the nucleus.

### AraC-p75^NTR^- mediated inactivation of NFκB pathway exacerbates neurodegeneration by activating cell death/JNK pathway

We previously reported that uncoupling of p75^NTR^ from NF*κ*B leads to cell death by activation of the JNK pathway^43^. In the absence of RIP2, an adaptor protein linking p75^NTR^ to the NF*κ*B pathway, p75^NTR^ binds to TRAF6, another adaptor protein that links p75^NTR^ to the JNK apoptotic pathway. For this reason, we asked whether AraC-mediated inactivation of NF*κ*B pathway could lead to the activation of the JNK pathway, exacerbating neuronal apoptosis. The recruitment of TRAF6 to the intracellular domain of p75^NTR^ after AraC treatment was assessed in the CGNs cultures at 4 DIV by proximity ligation assay (PLA). We detected an increase in p75^NTR^:TRAF6 PLA puncta (Fig 7a-b) that suggested that AraC alters the conformation of p75^NTR^ favouring the binding of TRAF6. Next, we evaluated the activation of the JNK apoptotic pathway by assessing phosphorylation of c-Jun on threonine 91 (Thr91), which has been linked to cell death in CGNs^59^. Indeed, binding of AraC to WT p75^NTR^ increased the phosphorylation of c-Jun on Threonine 91 residue (Fig 7c-d). All together, these data indicate that treatment of CGNs with AraC inhibits the NF*κ*B survival pathway and potentiates the JNK apoptotic pathway.

**Figure 7.**
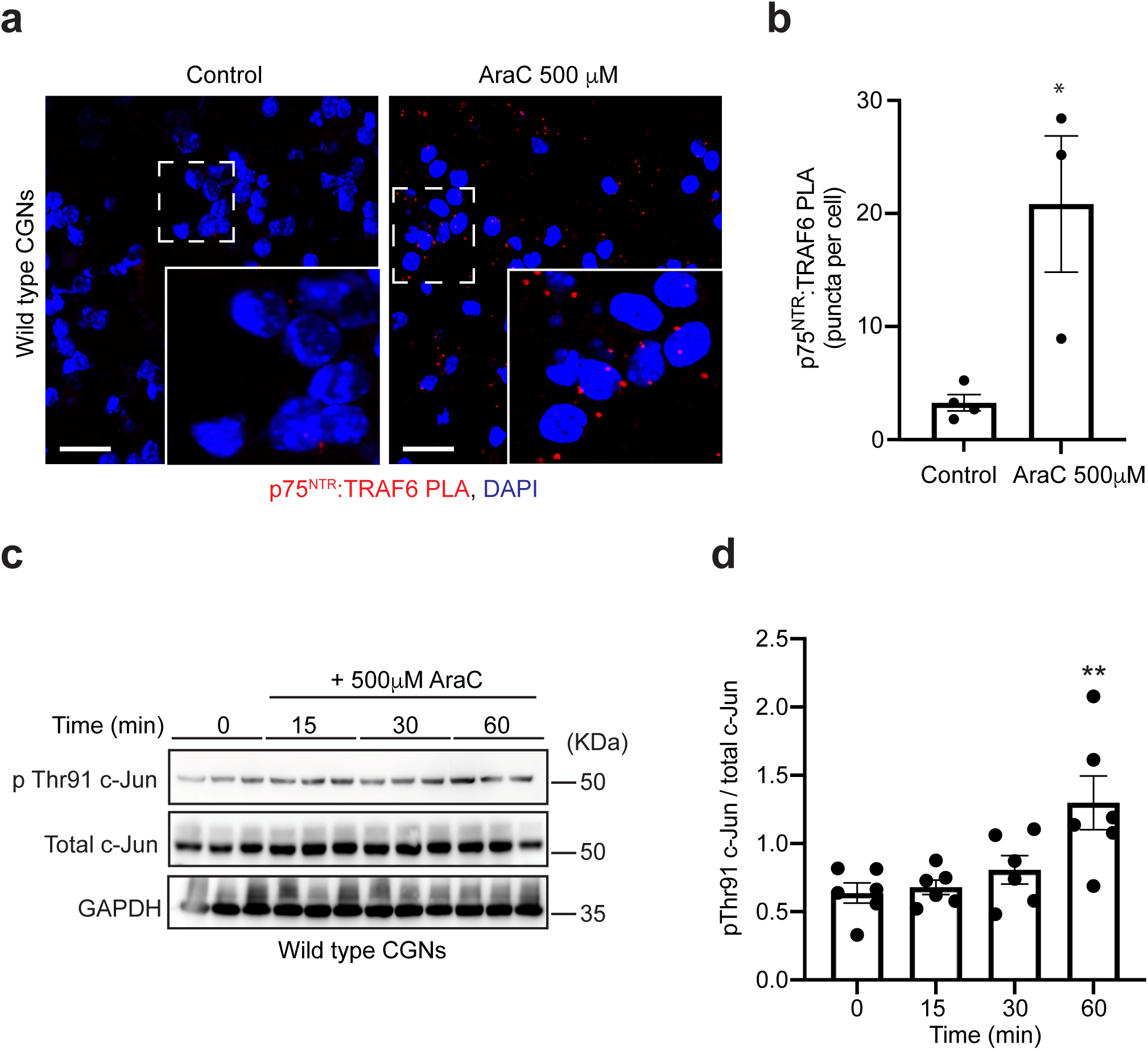
AraC/p75^NTR^- mediated inactivation of NFκB pathway exacerbates neuronal death by activating JNK pathway. (**A**) Micrographs of p75^NTR^:TRAF6 PLA (red) in untreated (control) or 500μM AraC treated CGNs for 10 minutes. Images were selected from 25 images per condition from 3-4 separate experiments. The scale bar represents 20 μm. (**B**) Quantification of p75^NTR^:TRAF6 PLA puncta in untreated (control) or 500μM AraC treated CGNs for 10 minutes. Mean ± sem of data from 3 experiments (*P < 0.05; Unpaired Student T-test) is shown. (**C**) Representative western blots probed with phospho-c-Jun (Thr91), total c-Jun and GAPDH of lysates of wild type P7 CGNs grown for 4 days prior to 15-, 30- or 60-minutes treatment with 500μM AraC. (**D**) Quantification of c-Jun (Thr91) phosphorylation of total lysate of untreated wild type P7 CGNs or neurons treated with 500μM for 15, 30 and 60 minutes. Mean ± sem of densitometry from 4 experiments (*P < 0.05; oneway ANOVA followed by Bonferroni test) is shown.

## Discussion

AraC has successfully been used as a chemotherapeutic for decades, being the most effective chemotherapy for the treatment of several cancers including AML, acute lymphatic leukaemia (ALL) and non-Hodgkin’s lymphoma^1–4^. Similar to other chemotherapy treatments, AraC has several side effects including neurotoxicity^3, 6, 7^. Interestingly, AraC-induced neurotoxicity is age-dependent, the older the patient is the more severe the neurotoxicity^11^. In the current study, we show that high doses of AraC induces apoptosis in mature cerebellar neurons. This results is in agreement with observations made in adult cancer patients under HIDAC treatment regime, who develop cerebellar atrophy and shrinkage leading to impaired cerebellar function that in some cases was permanent^11, 36^. As expected, the mature CGNs *in vitro* express the neurotrophic receptor, p75^NTR^. Although, it has been suggested that p75^NTR^ expression is markedly reduced in adult neurons^17^, the expression of this receptor is not completely abrogated in adult cerebellum^19, 43^. Interestingly, acute leukaemia patients show expression of p75^NTR^ both on malignant and normal lymphocytes, as well as an increase in expression of p75^NTR^ in serum, bone marrow and peripheral blood cells ^24, 25^. Therefore, it is plausible that there is also an increase of p75^NTR^ in the cerebellum of these patients making them more susceptible to AraC mediated neurodegeneration through p75^NTR^. In agreement with this, our data demonstrates that deletion of p75^NTR^ in matured CGNs prevented AraC-induced apoptosis.

P75^NTR^ facilitates different effects in CGNs, including axonal degeneration, cell survival, apoptosis and growth inhibition ^22, 23^. These effects are achieved due to p75^NTR^ capability to couple different signalling pathways including NF*κ*B, JNK/caspase and RhoDGI/RhoA pathways ^23, 53^ in these neurons. This raises the question of why the binding of AraC to p75^NTR^ induces neurite degeneration and cell death and not survival or growth inhibition in mature CGNs. It is noteworthy that the outcome of AraC binding to p75^NTR^ depends on the availability of adaptor proteins and the signalling pathway that will be engaged.

Surprisingly, we find that AraC binds to the TMD of p75^NTR^. Recent findings have demonstrated that a couple of small molecules binding to TMD of receptors, including SB394725 that binds TMD of thrombopoetein receptor^60^ and NCS49652, a compound that binds to the TMD of p75^NTR 33^. The binding of AraC to p75^NTR^ TMD leads to anisotropy oscillations that are different to those induced by the binding of NSC49652^33^, suggesting that the two compounds binds to different regions of the p75^NTR^ TMD. In fact, while NSC496532 binds to Ile252, Pro253, Val 254, Cys256, and serine (Ser) 257^33^, our data suggest that the binding of AraC to p75^NTR^ TMD only require Cys256 and Pro253.We suggest that this specificity and the positions of these two residues could be the reason why AraC does not bind to other members of the TNF superfamily that p75^NTR^ belongs to^61, 62^.

Interestingly, the binding of AraC to Cys256 and Pro253 residues in p75^NTR^ TMD enables AraC to specifically uncouple p75^NTR^ from the NF*κ*B pathway and not any other p75^NTR^- dependent pathway. We previously reported that uncoupling of p75^NTR^ from the NF*κ*B pathway leads to apoptosis of CGNs during cerebellar development^43^. Moreover, we showed that in the absence of RIP2, p75^NTR^ recruits TRAF6, which couples the receptor to the JNK pathway, leading to an increase in the apoptotic activity of CGNs^43^. Although the wild type neurons in the current study express both RIP2 and TRAF6, we speculated that the binding of AraC to p75^NTR^ TMD changes the conformation of p75^NTR^ hindering the recruitment of RIP2 to p75^NTR^ death domain. This new conformation favours the binding of TRAF6 to p75^NTR^ inducing the apoptotic activity of p75^NTR^ observed in these neurons. Indeed, our data shows that treatment of mature CGNs with AraC leads to increased p75^NTR^:TRAF6 interaction rendering its specificity of activating the TRAF6/JNK cell death pathway.

Although the current work has focused on the effect of AraC on mature CGNs, we speculate that mature neurons in other brain regions that express p75^NTR^ could be also affected by this treatment. Interestingly, AraC treated patients also exhibit somnolence and drowsiness^63, 64^, reinforcing the notion that this drug could alter the function of other brain regions besides the cerebellum. Intriguingly, one brain region that express p75^NTR^ in abundance in adult brain is the basal forebrain^18^, which has been implicated in the homeostatic regulation of sleep^65–67^. It is therefore plausible that the somnolence and drowsiness phenotype observed in patients on HIDAC is a result of AraC/p75^NTR^-mediated neurite degeneration and cell death of the neurons in basal forebrain.

### Conclusions

In conclusion, our data show that p75^NTR^ facilitates AraC-induced degeneration of mature CGNs by uncoupling p75^NTR^ from NF*κ*B pathway and exacerbating cell death/JNK pathway, contributing to cerebellar degeneration. Our data elucidates the molecular mechanisms of action of the AraC in the degeneration and cell death of mature neurons, providing a new molecular target to develop treatments to counteract the side effects of AraC in the CNS.

## Abbreviations

AKT: protein kinase B
ALL: acute lymphatic leukaemia
AML: acute myeloid leukaemia
ANOVA: analysis of variance
AraC: 1-beta-D-arabinofuranosyl-cytosine cytarabine;
AraTM: ara transmembrane;
BCA: bicinchoninic acid assay
cDNA: complementary DNA
CETSA: the cellular thermal shift assay,
CETSA: the cellular thermal shift assay,
CGNs: cerebellar granule neurons;
CNS: centran nervous system
COS-7: -African green monkey kidney fibroblast-like cell line.
Cys: cystein
DI: degeneration index
DIV: days in vitro
DMSO: dimethyl sufoxide
DNA: deoxyribonucleic acid
ECL: enhanced chemiluminescence
EGFP: enhanced green fluorescent protein
FRET: förster Resonance Energy Transfer;
GAPDH: glyceraldehyde 3-phosphate dehydrogenase
HEK: human embryonic kidney cells
HEPES: 4-(2-hydroxyethyl1)-1-piperazineethanesulfonic acid
HIDAC: high Dose AraC
HRP: horseradish peroxidase
IgG: immunoglobulin
IkBa: nuclear factor of kappa light polypeptide gene enhancer in B-cells inhibitor, alpha
Ile: isoleucin
IPTG: isopropyl B-D-1-thiogalactopyranoside
ITDR-CETSA: Isothermal dose response-cellular thermal shift assay
JNK: c-Jun N-terminal kinase
KCL: potassium chloride
LB: lysogeny broth
NFkB: nuclear factor kappa-light-chain-enhancer of activated B cells
NIH: national institute of health
NTR: neurotrophine receptor
PBS: phosphate buffered solution
PCR: polymeras chain reaction
PLA: *Proximity Ligation Assay*
RhoA: ras homolog family member A
RhoGDI: Rho GDP-dissociation inhibitor
RT: room temperature
Ser: serin
TAG-1: transiet axonal glycoprotein
TMD: transmembrane domain
TNF: trumor necrosis factor
TRAF6: TNF receptor associated factor 6
TrkB: Tropomysin receptor kinase B
TUNEL: terminal deoxynucleotidyl transferase dUTP nick end labelling
Val: valin
WT: wild type

## Declarations

### Ethical approval

All animal experiments were conducted in accordance with the National University of Singapore Institutional Animal Care, Use Committee and the Stockholm North Ethical Committee for Animal Research regulations and the Danish Animal Experiment Inspectorate Under the Ministry of Justice.

### Consent of publication

Not applicable

### Availability of data and materials

All the data used and analyzed for this study are available from the corresponding author on reasonable request.

### Competing financial interest

The author declares no competing financial interests.

### Funding

This research was supported by grants to LK from Karolinska Institute Research Foundation and Magnus Bergwall Foundation and to AN from The Danish National Research Foundation grant DNRF133

### Author contributions

VLR performed TM binding assay and Homo-FRET experiments. PB performed neurite degeneration and PLA experiments. DIP and LK performed the CESTA experiment. JCR, AA and DFS performed initial experiments. AN provided reagents and funding. LK conceived the study, planned, performed and analysed the majority of the experiments. LK wrote the manuscript with input from all authors. All authors read and approved the final manuscript.

## Acknowledgments

We thank professor Carlos F Ibáñez, Karolinska Institute for providing resources and insightful discussion. We also thank Dr. Stella Nolte, Anne-Kerstine Glintborg Jensen, Andreea-Cornelia Udrea and Ket Yin Goh for excellent technical support.

## Author’s information

**Vanessa Lopes-Rodrigues**: Department of Physiology and Life Sciences Institute, National University of Singapore, Singapore 117597, Singapore. **Email**

**Pia Boxy:** Danish Research Institute of Translational Neuroscience (DANDRITE)–Nordic EMBL Partnership for Molecular Medicine, Department of Biomedicine, Aarhus University, Denmark. The Danish National Research Foundation Center, PROMEMO, Department of Biomedicine, Aarhus University, Aarhus, Denmark. **Email:**

**Eunice Sim:** Department of Physiology and Life Sciences Institute, National University of Singapore, Singapore 117597, Singapore. **Email:**

**Dong Ik Park:** Danish Research Institute of Translational Neuroscience (DANDRITE)–Nordic EMBL Partnership for Molecular Medicine, Department of Biomedicine, Aarhus University, Denmark. The Danish National Research Foundation Center, PROMEMO, Department of Biomedicine, Aarhus University, Aarhus, Denmark. **Email:**

**Josep Carbonell:** Department of Neuroscience, Karolinska Institute, Stockholm S-17177, Sweden. **Email:**

**Annika Andersson:** Department of Neuroscience, Karolinska Institute, Stockholm S-17177, Sweden. **Email:**

**Diana Fernández-Suárez:** Department of Neuroscience, Karolinska Institute, Stockholm S-17177, Sweden. **Email:**

**Anders Nykjær:** Danish Research Institute of Translational Neuroscience (DANDRITE)– Nordic EMBL Partnership for Molecular Medicine, Department of Biomedicine, Aarhus University, Denmark. The Danish National Research Foundation Center, PROMEMO, Department of Biomedicine, Aarhus University, Aarhus, Denmark. **Email:**

**Lilian Kisiswa:** Danish Research Institute of Translational Neuroscience (DANDRITE)– Nordic EMBL Partnership for Molecular Medicine, Department of Biomedicine, Aarhus University, Denmark. The Danish National Research Foundation Center, PROMEMO, Department of Biomedicine, Aarhus University, Aarhus, Denmark. **Email:**

## Notes

### Competing Interest Statement

The authors have declared no competing interest.

